# CRISPR-Cas systems restrict horizontal gene transfer in *Pseudomonas aeruginosa*

**DOI:** 10.1101/2020.09.19.304717

**Authors:** Rachel M. Wheatley, R. Craig MacLean

## Abstract

CRISPR-Cas systems provide bacteria and archaea with an adaptive immune system that targets foreign DNA. However, the xenogenic nature of immunity provided by CRISPR-Cas raises the possibility that these systems may constrain horizontal gene transfer. Here we test this hypothesis in the opportunistic pathogen *Pseudomonas aeruginosa*, which has emerged an important model system for understanding CRISPR-Cas function. Across the diversity of *P. aeruginosa*, active CRISPR-Cas systems are associated with smaller genomes and a reduced GC content, suggesting that CRISPR-Cas inhibits the acquisition of foreign DNA. Although phage are the major target of CRISPR-Cas spacers, more than 80% of isolates with an active CRISPR-Cas system have spacers that target integrative conjugative elements (ICE) or the conserved conjugative transfer machinery used by plasmids and ICE. Consistent with these results, genomes containing active CRISPR-Cas systems harbor a lower abundance of both prophage and ICE. Crucially, spacers in genomes with active CRISPR-Cas systems map to ICE and phage that are integrated into the chromosomes of closely related genomes lacking CRISPR-Cas immunity, providing direct evidence that CRISPR-Cas constrains horizontal gene transfer in these lineages. In conclusion, we find that CRISPR-Cas acts as an important constraint to horizontal gene transfer, suggesting that CRISPR-Cas may constrain the ability of this pathogen to adapt to new niches and stressors.

## Introduction

CRISPR (clustered regularly interspaced short palindromic repeats)-Cas(CRISPR associated protein) systems are adaptive immune systems that provide heritable immunity against foreign DNA, and are widespread in bacterial and archaeal genomes (1-3). CRISPR-Cas systems are able to incorporate segments of invading DNA, such as may come from bacteriophage or mobile genetic elements, as spacers in CRISPR loci (4). Active systems must contain a set of Cas genes that enable the CRISPR arrays to be transcribed and processed into short CRISPR RNAs (crRNAs) (5). These crRNAs contain a single spacer and must be bound to the Cas endonuclease. This complex uses crRNA base complementarity to recognise and degrade DNA from elements containing the spacer sequence upon subsequent re-infection of the cell (6). In effect, CRISPR-Cas systems provide a molecular memory of past infections and provide bacteria and archaea with adaptive immunity against foreign DNA (1, 4, 6).

Horizontal gene transfer (HGT) plays an important role in bacterial evolution (7), and is a major source of genome expansion (8). CRISPR-Cas systems were first recognised for their role as phage defence mechanisms (3, 9-11), and can provide protection by preventing lysogenic conversion (12, 13), which is an important mechanism of HGT (14-16). Although CRISPR-Cas systems can target parasitic genetic elements such as lytic phage, the xenogenic immunity provided by CRISPR-Cas may constrain HGT more broadly (17-20). There is growing recognition that CRISPR-Cas systems target other mobile genetic elements (17, 21, 22), and it is suggested that CRISPR-Cas may play a very general role in preventing HGT by targeting integrative conjugative elements (ICE) and plasmids, or DNA that is acquired by transformation. Experimental studies have shown in a number of systems that CRISPR-Cas can prevent HGT over short time scales (3, 11, 21, 23). For example, in *Staphylococcus epidermidis*, CRISPR-Cas systems possessing a spacer that targets a highly conserved nickase present on staphylococcal conjugative plasmids have been shown to be successful in preventing plasmid transformation (21). Bioinformatic studies, on the other hand, have produced conflicting results on the importance of CRISPR-Cas in HGT over longer time scales (18, 21, 23-27). For example, a recent bioinformatics study found no evidence of a correlation between CRISPR-Cas activity and the frequency of HGT (24). Furthermore, a genome-wide correlation analysis reported that the presence of CRISPR-Cas systems constrains the acquisition of antibiotic resistance genes in only a sub-set of bacterial pathogens (18). In summary, the role of CRISPR-Cas is well established from an experimental point of view, but the long-term consequences of this interference are not as clearly understood. In this paper, we tackle this problem by testing the role of CRISPR-Cas systems on HGT in the opportunistic pathogen *Pseudomonas aeruginosa.*

*P. aeruginosa* genomes are large (typically 6-7 Mbp), and ∼50% of sequenced *P. aeruginosa* genomes have been predicted to possess an active CRISPR-Cas system (18, 28). Three major CRISPR-Cas system types (I-F, I-E and I-C) have been identified (28), and *P. aeruginosa* genomes contain a large repertoire of mobile genetic elements, including phage, transposons, ICE and plasmids (29, 30). ICE are modular mobile genetic elements that can integrate into a host genome and be vertically propagated through cell replication or transfer horizontally following excision from the chromosome (31, 32). ICE and plasmids both use the same type IV secretion system for conjugative transfer (31, 33-35), and the difference between ICE and plasmids comes from their ability to integrate into the chromosome. Like plasmids, ICE contain cargo genes (29, 36-38), and *P. aeruginosa* ICE have been implicated in a range of traits including xenobiotic compound degeneration (39), antibiotic resistance (32, 40-42), and virulence formation (43). Although plasmids and ICE share many similarities, ICE are abundant in *P. aeruginosa*, whereas plasmids are thought to be comparatively rare.

While it is straightforward to understand the benefits of CRISPR based immunity to obligate genetic parasites, such as lytic phage. Mobile genetic elements, on the other hand, can be either parasitic or beneficial, depending on conditions. For example, the acquisition of prophage can alter improve *Pseudomonas* metabolism and increase competitive ability (16, 44), but prophage entry into the lytic cycle leads to cell lysis and death. Similarly, ICE and plasmids carry genes that can allow *Pseudomonas* to exploit new niches, such as novel metabolites or eukaryotic hosts, or resist stresses, such as heavy metals and antibiotics, but the acquisition of these elements also tends to be associated with costs that can generate selection against carriage (32, 45-47). Given these costs and benefits, it is difficult to predict whether CRISPR-Cas systems should target these elements. The diversity and plasticity of *P. aeruginosa* genomes combined with the high variability of CRISPR-Cas presence makes *P. aeruginosa* a very useful species to study for evidence of CRISPR-Cas mediated inhibition of HGT. Previous work has shown that *P. aeruginosa* CRISPR-Cas systems are associated with small genome size (28), and reduced mobile sulphonamide resistance genes (18), and it has been suggested that *P. aeruginosa* is an example of a bacterial pathogen where CRISPR-Cas does play a recognisable role in HGT. However, the broader impacts of CRISPR-Cas on HGT and genome divergence have not been investigated in this species in detail. Here we analyse 300 high-quality assembled genomes (including 201 complete genomes) of *P. aeruginosa* to test the hypothesis that CRISPR-Cas constrains HGT, and to identify mobile genetic elements that are targeted by CRISPR-Cas. There is growing evidence that anti-CRISPR (*acr*) genes play an important role in antagonizing CRISPR-Cas (48-51), and our analysis also tests the hypothesis that *acr* genes negate the impact of CRISPR-Cas on HGT.

## Materials and Methods

### Genomic data

*P. aeruginosa* genome sequences were downloaded from NCBI RefSeq(https://ftp.ncbi.nlm.nih.gov/genomes/refseq/bacteria/Pseudomonas_aeruginosa/) (*Supplementary Information Table 1*). These genomes are considered complete or assembled to a high level; complete genome (201), chromosome (39) or scaffolds of 7 or fewer (60). Genome metadata was downloaded in parallel (genome size, guanine-cytosine (GC) content, number of coding sequences (CDS)). Multi-locus sequence typing (MLST) was carried out using MLST software developed that scans against PubMLST typing schemes (https://pubmlst.org/) (52, 53). Genome annotation was carried out using prokka (54).

**Table 1.**
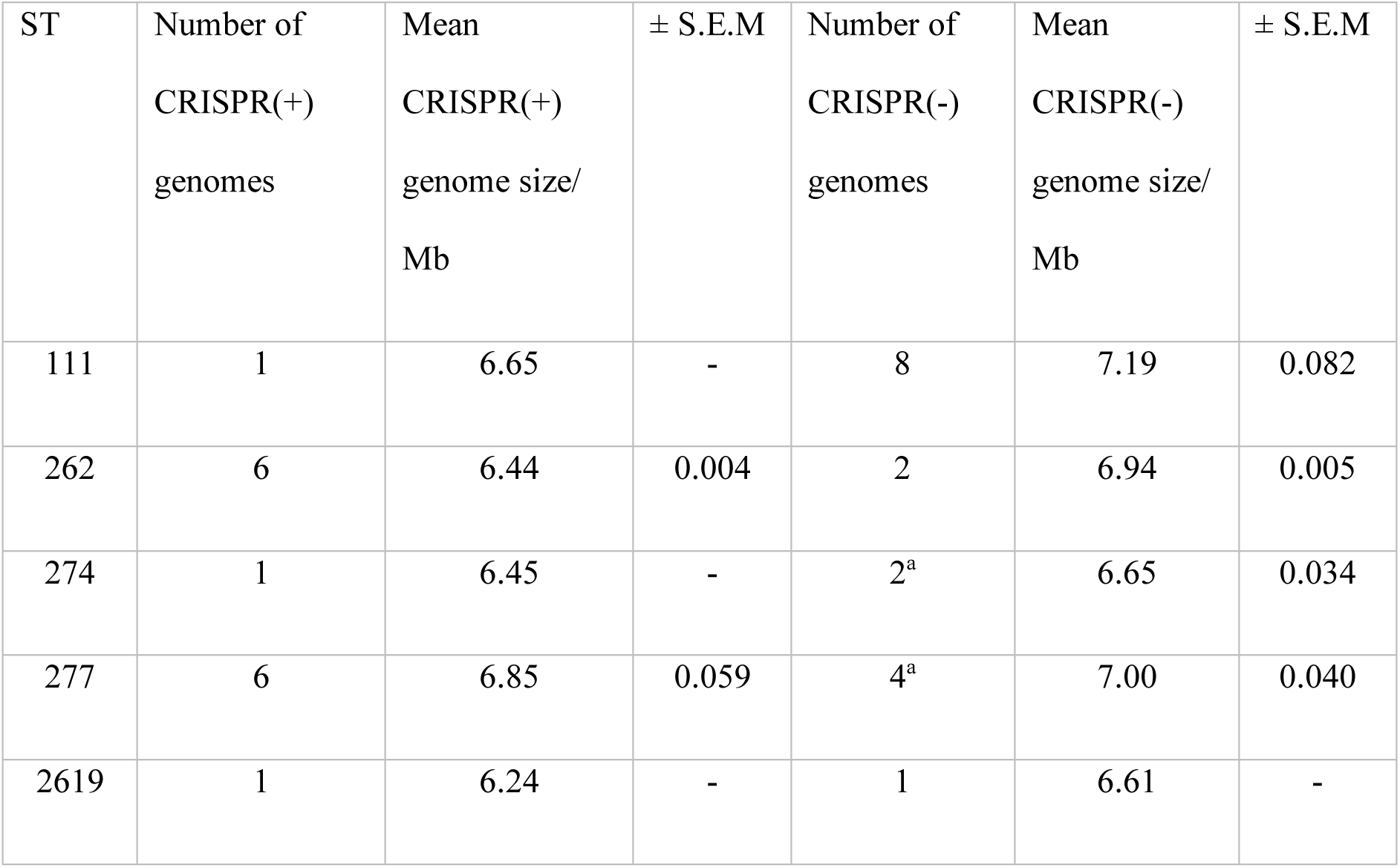
*P. aeruginosa* STs with variable presence of CRISPR-Cas systems: ST111, ST262, ST274, ST277 and ST2619. Table shows the number of CRISPR(+) and CRISPR(-) genomes in each ST, and the size (Mb) of these genomes (mean if > 1 genome). CRISPR(-) genomes were larger than their CRISPR(+) counterparts (paired sample t-test p < 0.01). It should be noted ST262 genomes are from the same bioproject (*Supplementary Information Table 2*). ^a^; one genome within this subset is CRISPR(-) by virtue of containing Acr.

| ST | Number of<br>CRISPR(+)<br>genomes | Mean<br>CRISPR(+)<br>genome size/<br>Mb | ± S.E.M | Number of<br>CRISPR(-)<br>genomes | Mean<br>CRISPR(-)<br>genome size/<br>Mb | ± S.E.M |
| --- | --- | --- | --- | --- | --- | --- |
| 111 | 1 | 6.65 | - | 8 | 7.19 | 0.082 |
| 262 | 6 | 6.44 | 0.004 | 2 | 6.94 | 0.005 |
| 274 | 1 | 6.45 | - | 2<sup>a</sup> | 6.65 | 0.034 |
| 277 | 6 | 6.85 | 0.059 | 4<sup>a</sup> | 7.00 | 0.040 |
| 2619 | 1 | 6.24 | - | 1 | 6.61 | - |

### CRISPR-Cas and anti-CRISPR annotation

CRISPRCasFinder was used to predict presence of CRISPR arrays and cognate Cas proteins (55). CRISPRCasFinder assigns evidence levels to putative CRISPR loci on a 1-4 level scale (55), using an algorithm to measure CRISPR repeat conservation based on Shannon’s entropy and produce an EBcons (entropy-based conservation) index. Evidence level 4 was used as the cut-off for annotating CRISPR loci (*Supplementary Information Table 1*), and the details of this algorithm and evidence level system are described in (55). CasFinder version 2.0 of CRISPRCasFinder was used to identify and type Cas systems in genomes with predicted CRISPR loci (*Supplementary Information Table 1*) (55, 56). Anti-CRISPR (Acr) genes were identified through screening (blastn, default settings (57, 58)) genomes against Acr sequences used by van Belkum *et al* (28) and additional *P. aeruginosa* Acr sequences in the anti-CRISPR database (51). CRISPR-Cas systems were predicted to be functionally active if they were annotated to possess a CRISPR array, cognate Cas genes and the absence of Acr. All Cas systems were identified to be type I-F, I-E or I-C, with the exception of one type U annotation (55), which was in a CRISPR(-) predicted genome (*Supplementary Information Table 1*) containing Acr.

### Spacer target identification

A unique spacer set (n = 2075) was generated by clustering spacer sequences identified in CRISPRCasFinder (55) for all CRISPR(+) genomes with CD-HIT (59, 60) using a 95% sequence identity threshold as used in previous studies (18). Blastn was used to predict spacer targets by screening unique spacers against five databases: 1) Phage genomes 2) ICE, plasmid and conjugative transfer gene sequences 3) Resistance genes 4) Virulence genes and 5) Coding sequences from *P. aeruginosa* genomes. Blastn hits with at least 95% sequence identity to a spacer and at least 95% sequence coverage were accepted as predicted spacer targets. This threshold was previously defined in a spacer analysis study by Shmakov *et al* (2017), based on control analysis against false positive predictions comparing prokaryotic to eukaryotic virus targeting (17). The phage genomes used in this study were downloaded from NCBI (ftp://ftp.ncbi.nih.gov/refseq/release/viral/), and the resulting database contained 12,182 genome sequences. The ICE, plasmid and conjugative transfer genes sequences were compiled from three locations. ICE sequences were downloaded from the ICEberg 2.0 database of bacterial integrative and conjugative elements (61). Plasmid sequences were downloaded from a curated database of plasmid sequences containing 10,892 complete plasmid sequences (62). Details of how this plasmid database has been curated are given in Brooks *et al* (2019) (62). Conjugative transfer gene sequences (*tra* genes, *trb* genes and type IV secretion system genes) were downloaded from annotated *P. aeruginosa* genes in NCBI gene (63). Acquired resistance gene sequences were downloaded from the ResFinder database of acquired antimicrobial resistance genes (64), and virulence genes were downloaded from the Virulence Factor Database (65, 66). A blastn database of coding sequences from *P. aeruginosa* genomes was compiled from clustering predicted coding sequences of *P. aeruginosa* genomes used in this study with CD-HIT using a 95% sequence identity threshold.

The spacers per genome were analysed for GC content and the prediction of phage or ICE and conjugative transfer system targeting. The focus of this downstream analysis are on ICE and phage, rather than plasmids. The abundance of ICE and prophage in *P. aeruginosa* genomes make them good targets to study with regards to CRISPR-Cas system correlations. Plasmids are thought to be comparatively rare, and the sample size of genomes in this study combined with the varied levels of assembly (∼2/3 complete genomes) provides a dataset that we believe is not well suited to assess correlations between plasmid presence and CRISPR-Cas. As such, plasmids have not been included in the spacers per genome or intra-ST variability analysis. However, an excellent recent bioinformatic study by O’Meara *et al* details a broad-scale analysis of plasmid carriage and CRISPR across bacterial species (19). The GC of each spacer was calculated using a Perl script available on GitHub (67). The average spacer GC per genome was then calculated using awk in command line [awk -v s=2 ‘NR<s{next} {c++; t+=$2} END{printf “%.2f (%d samples)\n”, t/c, c}’ outputfile_gc.txt]. The spacer sequences per genome were searched against the phage, ICE and conjugative transfer gene datasets previously outlined using blastn. Blastn hits with at least 95% sequence identity to a spacer and at least 95% sequence coverage were accepted as predicted spacer targets (18).

### Analysis of intra-ST CRISPR variability

Five *P. aeruginosa* STs were identified with variable presence or absence of CRISPR-Cas systems: ST111, ST262, ST274, ST277 and ST2619. Complete CRISPR(+) and CRISPR(-) genome representatives of ST111, ST262, ST277 and ST2619 were aligned in Mauve (progressive Mauve alignment with default settings) (68) using the GenBank (.gb) files downloaded from NCBI. The complete genome sequences used for alignment are indicated in *Supplementary Information Table 2*. Unique regions annotated to contain phage or ICE in the GenBank annotation file were highlighted on the Mauve alignments. To quantify the influence of phage and ICE, we systematically searched for phage and ICE in our genomes with intra-ST CRISPR variability. The identification of prophage regions was carried out using PHASTER (69), which was used to predict a total number of prophage regions within each genome, combined across all three region scoring levels (69) (*Supplementary Information Table 2*). The estimation of ICE abundance was carried out using a blastn search of the coding sequence annotation files (.ffn) obtained from prokka (54) against the database of ICE sequences (61) and conjugative transfer system genes in *P. aeruginosa*. From this, the number of coding sequences predicted to represent these conjugative elements within each genome was standardised per genome Mb. A genome size standardised measure of ICE abundance was used so that conjugative element integration could be compared regardless of the already apparent genome size bias between CRISPR(-) and CRISPR(+) isolates.

**Table 2.** The number of CRISPR(+) spacers predicted to target unique regions present in the CRISPR (-) genomes for the complete genome representatives of ST111, ST262, ST277 and ST2619 that were aligned (Fig. 5) (*Supplementary Information Table 2*). The predicted targets of the spacers is taken from the annotation of each genome. ^b^; targets annotated as hypothetical proteins whose identity was further investigated using NCBI blast to search for characterised homologous coding regions (>90% identity across whole length).

| Sequence Type | Total number of spacers | Number of CRISPR(+) spacers found in blastn of CRISPR(-) unique genomic regions | Predicted targeting identity of spacers (given as putative gene encoded product) |
| --- | --- | --- | --- |
| ST111 | 27 | 4 | 1. Intergenic region<br>2. Restriction endonuclease-like protein<sup>b</sup><br>3. Conjugative transfer system protein<br>4. Conjugative transfer system protein |
| ST262 | 42 | 7 | 1. Phage protein (possible phage transcriptional regulator)<sup>b</sup><br>2. Hypothetical protein<br>3. Integrating conjugative element protein<br>4. Hypothetical protein<br>5. Two binding sites of intergenic regions<br>6. Phage protein (hypothetical protein)<sup>b</sup><br>7. Regulatory/ putative secretion system component |
| ST277 | 41 | 8 | 1. Hypothetical protein<br>2. Intergenic region<br>3. Hypothetical protein<br>4. Phage protein (virion structural protein)<sup>b</sup><br>5. Phage protein (structural protein)<sup>b</sup><br>6. Phage protein (hypothetical protein)<sup>b</sup><br>7. Hypothetical protein<br>8. DUF-domain containing protein |
| ST2619 | 38 | 4 | 1. DUF-domain containing protein<br>2. S49 family peptidase |
|  |  |  | 3. Conjugative plasmid adhesion PilV |
|  |  |  | 4. DUF-domain containing protein |

Unique CRISPR(-) regions identified from Mauve alignment were extracted in nucleotide (.fasta) format. These were compiled to produce a single (.fasta) file containing all unique regions present in each CRISPR(-) genome compared to their CRISPR(+) counterpart within an ST (*Supplementary Information Table 2*). Blastn was used to predict whether the CRISPR(+) spacer sequences had targeting identity to these unique CRISPR(-) regions. Blastn hits with a minimum e value of 0.01 were accepted as predicted spacer targets (18). Putative identity of the predicted spacer targets was taken from the annotated .gb file for each genome. The identity of hypothetical protein encoding gene targets was further characterised using NCBI blast, searching for coding regions with high homology to the target (>90% identity across whole length). Targets that have been further characterised in this way have been indicated (^b^) in Table 2.

### Statistical analysis

Statistical testing was done using built-in methods in R (t.test, cor.test) (70). An unpaired two tailed t-test was used to test the association between CRISPR-Cas system presence and genome size, and CRISPR-Cas system presence and GC content. A paired sample two tailed t-test was used to test the association between genome GC and spacer GC. Pearson’s correlation coefficient test was used to test the correlation between phage targeting spacers and total spacers within a genome. For intra-ST analyses, differences between CRISPR(-) and CRISPR(+) genomes were analysed using a paired sample one tailed t-test.

## Results and discussion

### Phylogenetic distribution of CRISPR-Cas in collection of *P. aeruginosa* genomes

A collection of 300 *P. aeruginosa* genomes were downloaded from NCBI RefSeq (71). These genomes spanned a large number of Sequence Type (ST)s: 113 defined STs (271 genomes), and 29 genomes had an undefined ST (*Supplementary Information Table 1*). A total of 150 of the 300 genomes were predicted to possess CRISPR arrays and accompanying Cas genes (55) (*Supplementary Information Table 1*). Degenerate systems were identified in 26 genomes that possessed a CRISPR array but lacked cognate Cas genes (*Supplementary Information Table 1*). The proportion of genomes with predicted CRISPR-Cas systems compared to those without is in line with previous studies that have analysed a larger dataset (> 600) of *P. aeruginosa* genomes (18, 28).

CRISPR-Cas systems may lose their effectiveness by the acquisition of anti-CRISPR (Acr) proteins. Acr proteins originate from phage genomes (49), and inhibit targeting by CRISPR-Cas systems through a variety of distinct mechanisms (12, 49, 50). Screening these genomes against an Acr database identified Acr genes in 27/150 genomes with a CRISPR-Cas system, leaving 123 genomes that were predicted to encode functionally active CRISPR-Cas systems (CRISPR(+) genomes). CRISPR(+) genomes spanned 54 defined STs, with 12 genomes of undefined ST. We also identified Acr genes in genomes lacking CRISPR loci and/or Cas genes (*Supplementary Information Table 1*). Interestingly, the presence of active CRISPR-Cas systems was variable in some STs, whereas other STs consisted entirely of either CRISPR(+) or CRISPR(-) genomes (*Supplementary Information Table 1*). For a more detailed analysis of the phylogenetic distribution of CRISPR-Cas systems in *P. aeruginosa*, an excellent study was carried out in 2015 by van Belkum *et al* (28).

### Relationship between CRISPR-Cas systems and genome size

HGT is the key source of genome expansion in bacteria (8); for example, approximately 99% of the genes in γ-proteobacteria (including *P. aeruginosa*) are predicted to be acquired by HGT (72). Given this tight link between gene acquisition and HGT, active CRISPR-Cas systems should be associated with smaller genomes if CRISPR-Cas constrains HGT. In agreement with previous work (28), we found that CRISPR-Cas systems were associated with smaller *P. aeruginosa* genome size (Fig. 1A) (28). To further test the hypothesis that CRISPR-Cas constrains gene acquisition, we compared how genome size varied between *P. aeruginosa* genomes with or without Acr genes. Genomes with both CRISPR-Cas systems and Acr genes (CRISPR(+)/Acr) were significantly larger than CRISPR(+) genomes with no Acr genes (Fig. 1B). As Acr proteins inhibit CRISPR systems, this difference in genome size supports the hypothesis that active CRISPR-Cas systems limit HGT. Genomes lacking CRISPR loci and/or Cas genes were also found to contain Acr genes (CRISPR(-)/Acr), and were slightly larger than genomes that were both CRISPR(-) and Acr negative (Fig. 1C). We speculate whether this might be representative of strains that have acquired a large number of lysogenic phage, leading to both Acr acquisition and genome expansion.

**Figure 1.**
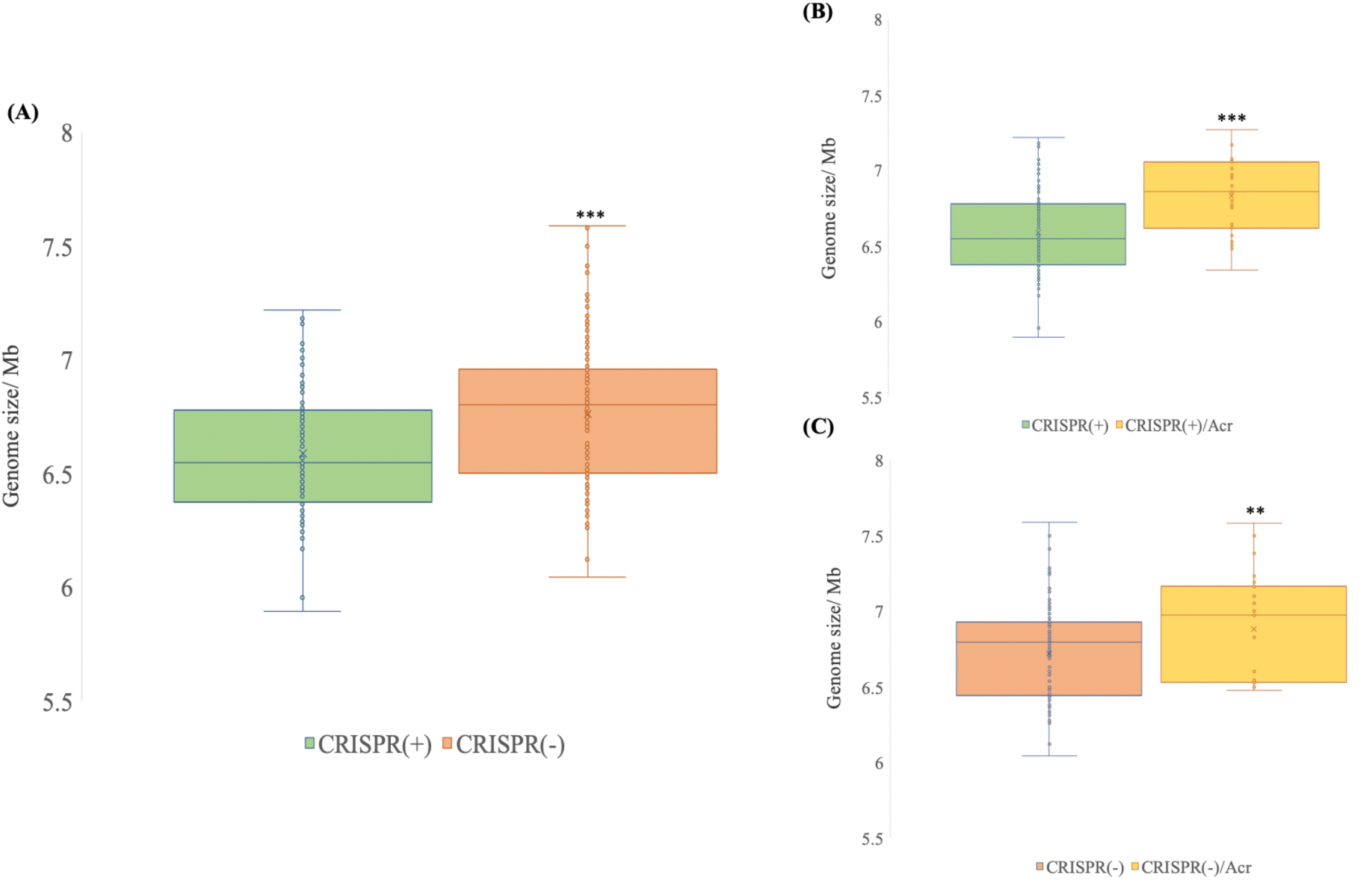
CRISPR and genome size. Panel A shows the size of CRISPR (+) and CRISPR (-) genomes; CRISPR (+) defined as genomes which contain a functionally predicted active CRISPR-Cas system and CRISPR (-) were defined as genomes lacking CRISRP-Cas and/or carrying Acr genes. The means of the two groups are significantly different at p < 0.001 (two tailed t-test). Panel B shows the size of genomes containing an active CRISPR-Cas system or both CRISPR-Cas and Acr genes. The means of the two groups are significantly different at p < 0.001 (two tailed t-test). Panel C shows the size of genomes that lack a CRISPR-Cas system that either carry or lack Acr genes. The means of the two groups are significantly different at p < 0.05 (two tailed t-test).

### Does CRISPR-Cas block the acquisition of potentially costly lower GC content elements?

Mobile genetic elements usually have a lower GC content than their bacterial hosts (73, 74), suggesting that HGT should be associated with reduced GC content. For example, the overall GC content of *P. aeruginosa* genomes is typically between 65-67% (29) which is very high in relation to most bacteria (75), and regions of the genome with low GC are indicative of the presence of recently acquired mobile elements (76, 77). We found that the presence of CRISPR-Cas was associated with higher genomic GC content, which is consistent with the hypothesis that CRISPR-Cas restricts the acquisition of foreign DNA (Fig. 2A). If CRISPR-Cas restricts the acquisition of mobile elements with low GC content, then we would also expect the GC content of CRISPR loci spacers to be low relative to the rest of the genome. Our findings support this; whilst genome GC stratifies by size, spacer GC was always lower than the genome-wide GC content, and the average difference in GC composition was 5% (Fig. 2B).

**Figure. 2.**
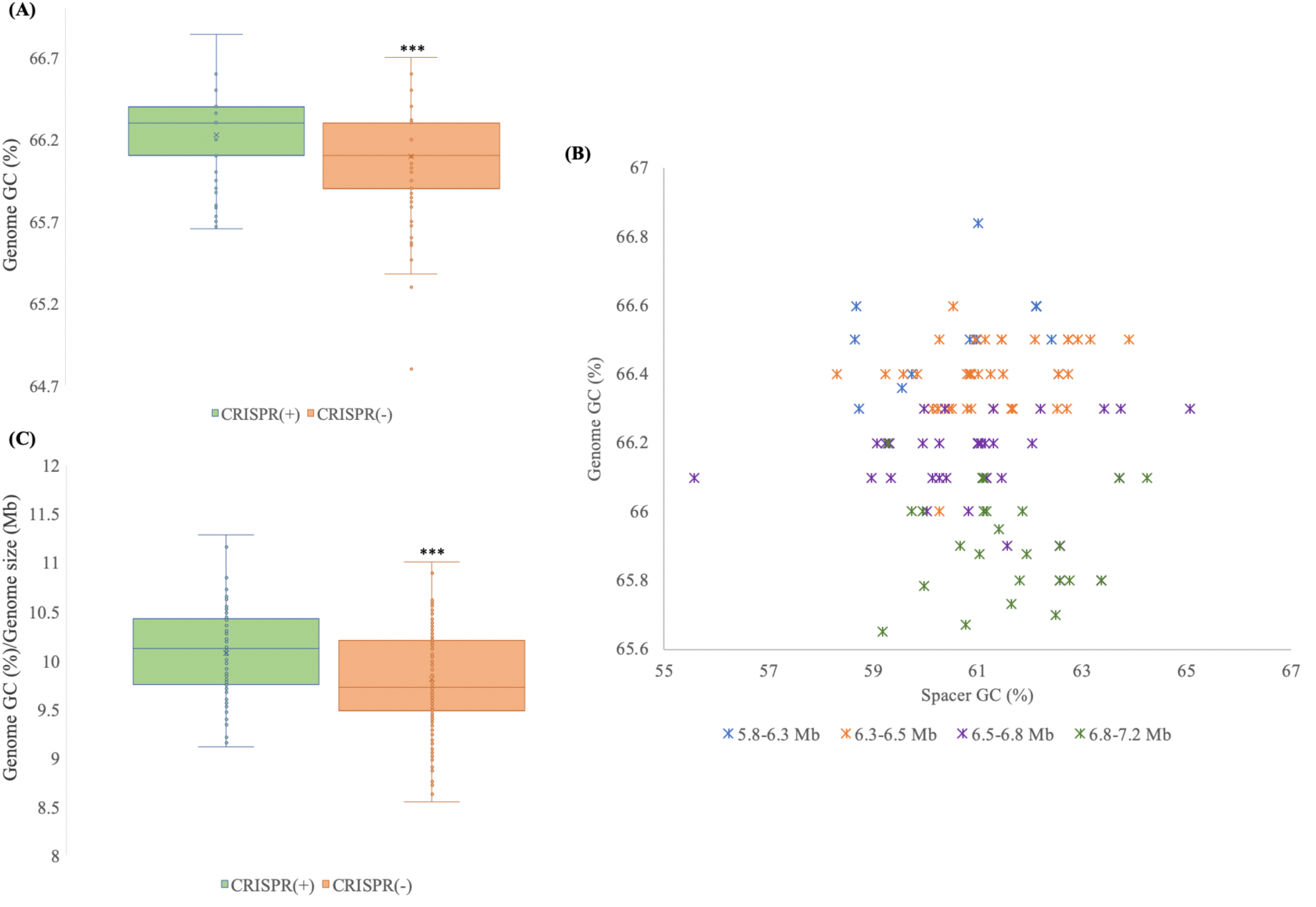
CRISPR-Cas systems and GC content. Panel A shows GC content in CRISPR(-) vs. CRISPR(+) genomes. Active CRISPR-Cas systems are associated with high GC content (two tailed t-test p < 0.001). Panel B shows a comparison of the GC content of spacers with genomic GC composition in genomes containing an active CRISPR-Cas system. Spacer GC content is always lower than genome-wide GC composition (paired sample t-test p < 0.001). Panel C shows the GC content of *Pseudomonas* genomes, standardized according to genome size. CRISPR-Cas systems are associated with higher genome GC after correcting for genome size (two tailed t-test p < 0.001).

Across bacterial species there is a correlation between genome size and GC content (78), suggesting that an association between functional CRISPR-Cas systems and GC content may be a spurious correlation driven by small size of CRISPR(+) genomes. However, we found that the association between CRIPSR-Cas activity and GC bias held true after correcting for variation in genome size (Fig. 2C). Although a complex number of factors can influence genome GC bias, our results suggest that CRISPR-Cas systems influence GC by preventing the acquisition of low GC elements.

### What are CRISPR-Cas loci spacers targeting?

To understand how CRISPR-Cas systems restrict HGT, we characterized the targets of CRISPR spacers in our collection. We identified a total of 2075 unique spacers across the CRISPR(+) *P. aeruginosa* genomes, and set out to characterise the proportion of this spacerome targeting phage, ICE, plasmids, and conjugative transfer genes, virulence factors, resistance genes and other coding sequences found in *P. aeruginosa* genomes (Fig. 3A). These screening categories were chosen to address key suggested targets of CRISPR-Cas systems (i.e. phage and mobile genetic elements) and potentially important genes involved in pathogenesis (i.e. resistance genes and virulence genes) that are also known to be carried on mobile genetic elements. Phage encounter is considered a strong evolutionary pressure for retaining CRISPR-Cas systems (12, 79) and, as expected, a large proportion of spacers (31.57%) were predicted to target phage DNA (Fig. 3A). Interestingly, the number of spacers targeting phage was positively well correlated to the total spacer number within each genome (Fig. 3B).

**Figure. 3.**
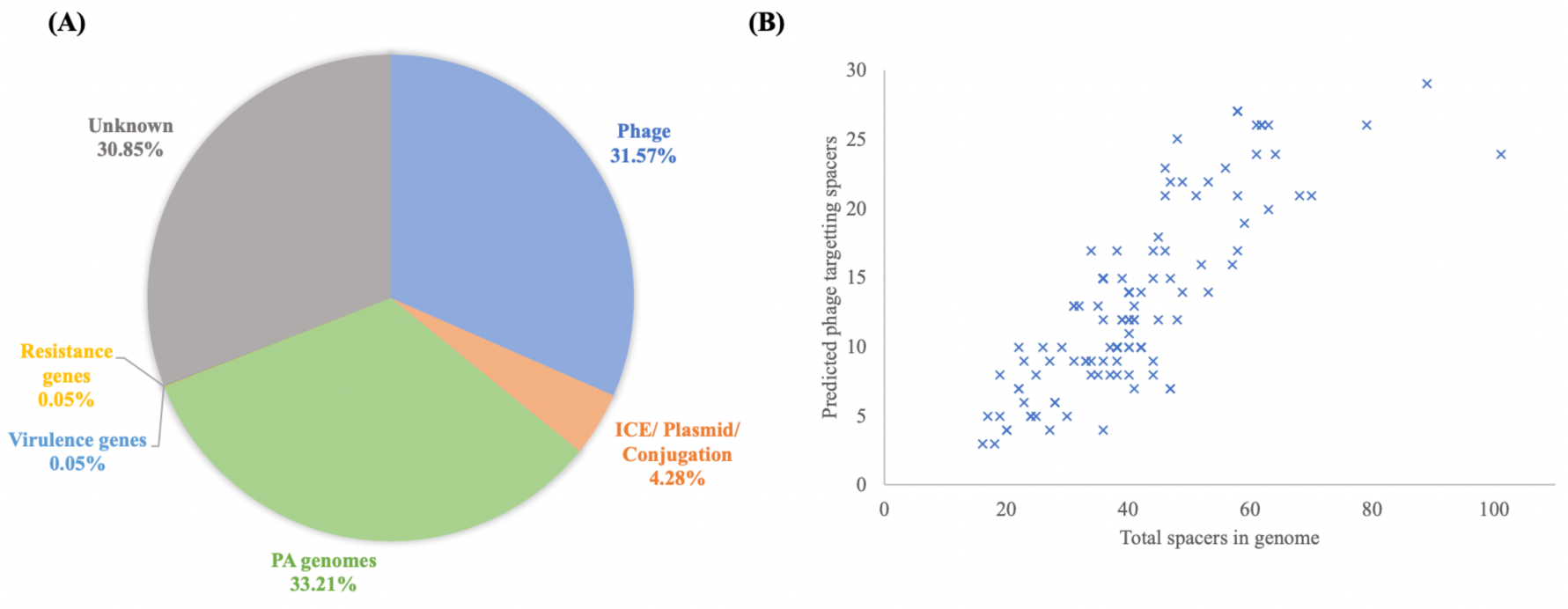
The CRISPR spacerome across *P. aeruginosa*. Panel A shows the predicted targets of the unique spacers in CRISPR(+) genomes across searches in databases for phage, ICE, plasmid, and conjugative genes, resistance genes, virulence genes and other coding sequences present in *P. aeruginosa* genomes. Panel B shows for each CRISPR(+) genome the total number of identified spacers and the number of these predicted to target phage. Predicted phage targeting spacers were strongly correlated to total genome spacer size (r=0.82, p<0.00001, Pearsons correlation coefficient test).

A smaller proportion of spacers (4.28%) were predicted to target ICE, plasmids and conjugative transfer genes (Fig. 3A). One unique spacer (0.05%) was predicted to target the *crpP* ciprofloxacin resistance gene (80), and this spacer was only found in one genome. Similarly, one unique spacer (0.05%) was predicted to target a virulence gene, which was a pyochelin dihydroaeruginoic acid synthetase gene (*pchE*), and this spacer was only found in one genome. Approximately 33% of spacers were predicted to target other coding sequences present in *P. aeruginosa* genomes (i.e. that do not overlap with characterised database sequences for phage, ICE, plasmids, and conjugative transfer genes, resistance genes or virulence factors). These were mainly annotated as coding sequences for hypothetical proteins. Approximately 31% of the spacers remained of unknown function (Fig. 3A), i.e., they had no identifiable target in our searches.

### Conjugative elements are a common target of *P. aeruginosa* CRISPR systems

We screened the spacers in each CRISPR(+) genome against a database of ICE and conjugative transfer genes to assess the prevalence of spacers targeting conjugative elements. Spacers targeting ICE or conjugative transfer system genes were widespread, occurring in 107/123 CRISPR(+) genomes (*Supplementary Information Table 1*), including 44/54 (∼81%) of the defined STs with an active CRISPR-Cas system in our collection (Fig. 4A). Crucially, spacers targeting conserved components of the conjugative machinery (*tra* genes, *trb* genes, type IV secretion system genes (81-83)) were found in in 25/54 (46%) of STs that were associated with active CRISPR-Cas systems (Fig. 4B). Although these spacers made up a small fraction of the spacerome, the distribution of these spacers and the high conservation of the conjugative machinery implies that CRISPR-Cas systems are likely to play an important role in preventing the acquisition of conjugative elements. In addition to these spacers, we identified spacers that target genes associated with ICE (61) in 42/54 (78%) of STs associated with active CRISPR-Cas systems (Fig. 4B). Ten STs contained no spacers predicted to target ICE or conjugative transfer system genes (Fig. 4A).

**Figure 4.**
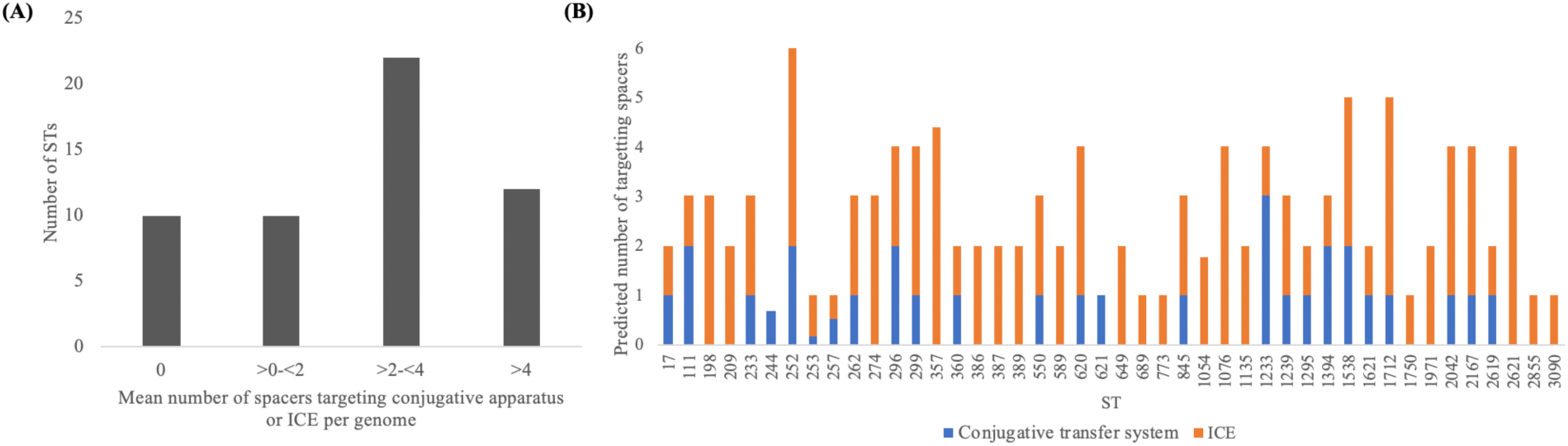
Spacers targeting ICE or conjugative transfer system genes. Panel A shows the combined number of ICE or conjugative transfer system targeting spacers per genome per ST across the 56 defined CRISPR(+) STs. Panel B shows the distribution of ICE and conjugative transfer system specific targeting spacers for the 44/54 defined STs they are present in, given as the mean per genome per ST.

### CRISPR-Cas systems prevent the acquisition of prophage and ICE

Our preceding analyses have focused on broad scale correlations between genome content and CRISPR-Cas activity across the diversity of *P. aeruginosa*. As a complementary approach, we focused in greater detail on five STs in our genome collection where the presence of predicted functional CRISPR-Cas systems was variable (Table 1). Since different isolates within the same ST are very closely related to each other, these STs provide the opportunity to investigate the evolutionary consequences of CRISPR-Cas activity in much greater detail, with less potential for confounding variables to obscure the effects of CRISPR-Cas. Consistent with our previous analyses, we found CRISPR(-) genomes were larger than their CRISPR(+) counterparts within an ST (Table 1).

We next investigated whether CRISPR-Cas systems could be directly linked to reduced gene acquisition in these STs. First, we aligned complete genome representatives of CRISPR(+) and CRISPR(-) from each ST, and then searched for unique regions containing phage or ICE, both of which make an important contribution to HGT in *Pseudomonas* (Fig. 5) (7, 37, 84). This alignment clearly suggested that the presence of CRISPR-Cas systems was associated with reduced occurrence of phage and ICE between pairs of closely related isolates (Fig. 5). To quantify this, we systematically searched for ICE and phage in all genomes in the five STs with variable presence or absence of CRISPR systems in our dataset (*Supplementary Information Table 2*).

**Figure 5.**
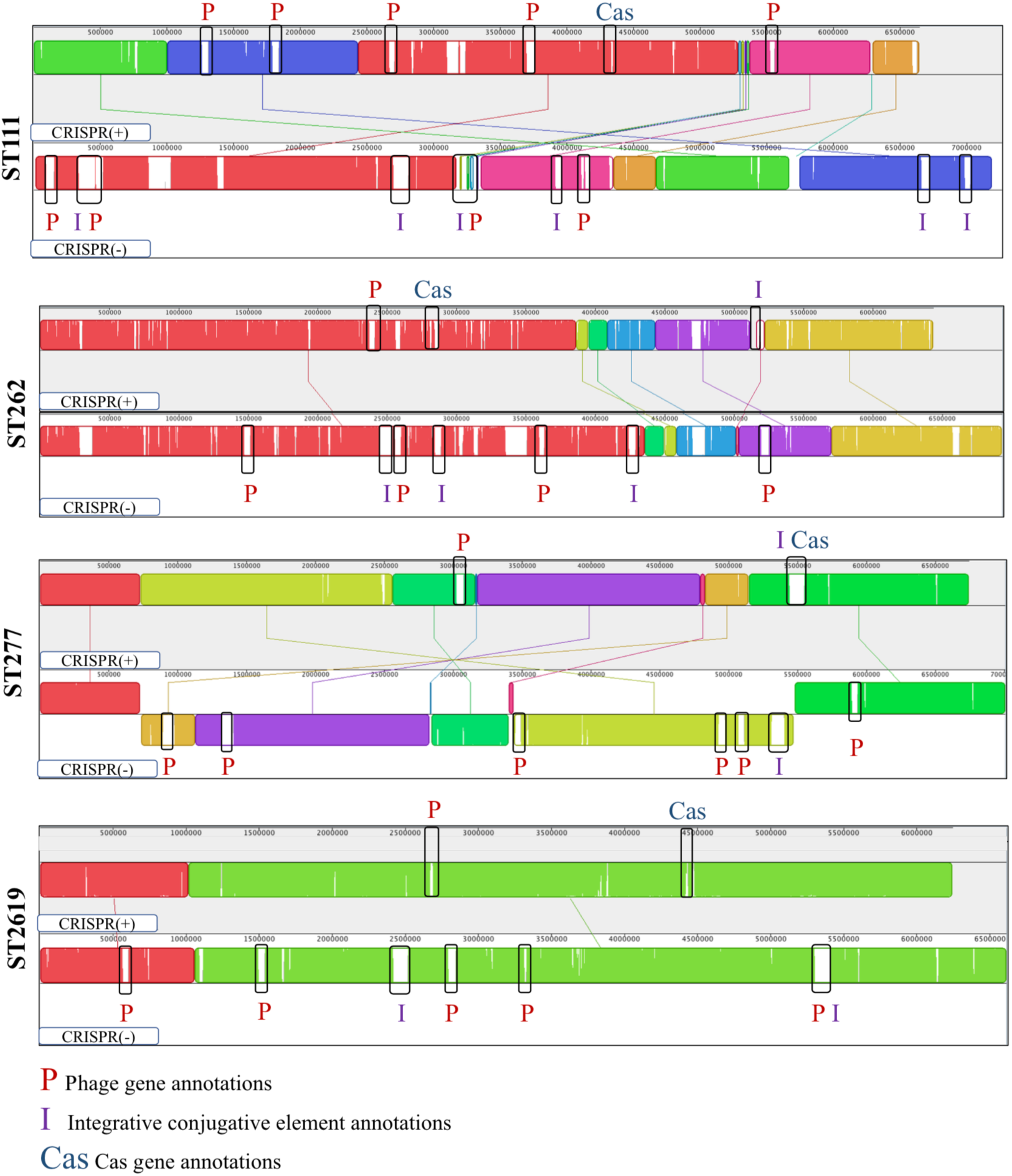
Mauve alignment of the complete CRISPR(+) and CRISPR(-) genome representatives of ST111, ST262, ST277 and ST2619 (68). Mauve alignment colours indicate colinear gene blocks. All eight complete genome representatives were composed of a chromosome and had no plasmids identified. Unique regions are shown by blank blocks of genes. Counterintuitively, a blank block of genes in a CRISPR(+) genome indicates genes that are absent from the corresponding CRISPR(-) genome, and vice versa. Unique regions were annotated according to the presence of phage (P), ICEs (I), or Cas genes (Cas). The genomes aligned are highlighted in Supplementary Table S2 and are complete genome sequences containing one chromosome. ST274 has been excluded from this alignment due to lack of a complete genome CRISPR(+) representative.

We found that functional CRISPR-Cas systems were associated with a lower relative abundance of predicted ICE genes in 4 of 5 STs, and the only exception (ST277) was one of the ten STs previously identified (Fig. 4A) that lacked any spacers predicted to target ICE or conjugative genes (Fig. 6A) (*Supplementary Information Table 2*). Similarly, we found that functional CRISPR-Cas systems were associated with a lower number of prophage regions (Fig. 6B) (*Supplementary Information Table 2*). Once again, ST277 was the exception to this general trend, and we speculate that CRISPR-Cas may have been recently gained or lost in this ST, given the small difference in genome size between CRISPR(+) and CRISPR(-) isolates in this ST.

**Figure 6.**
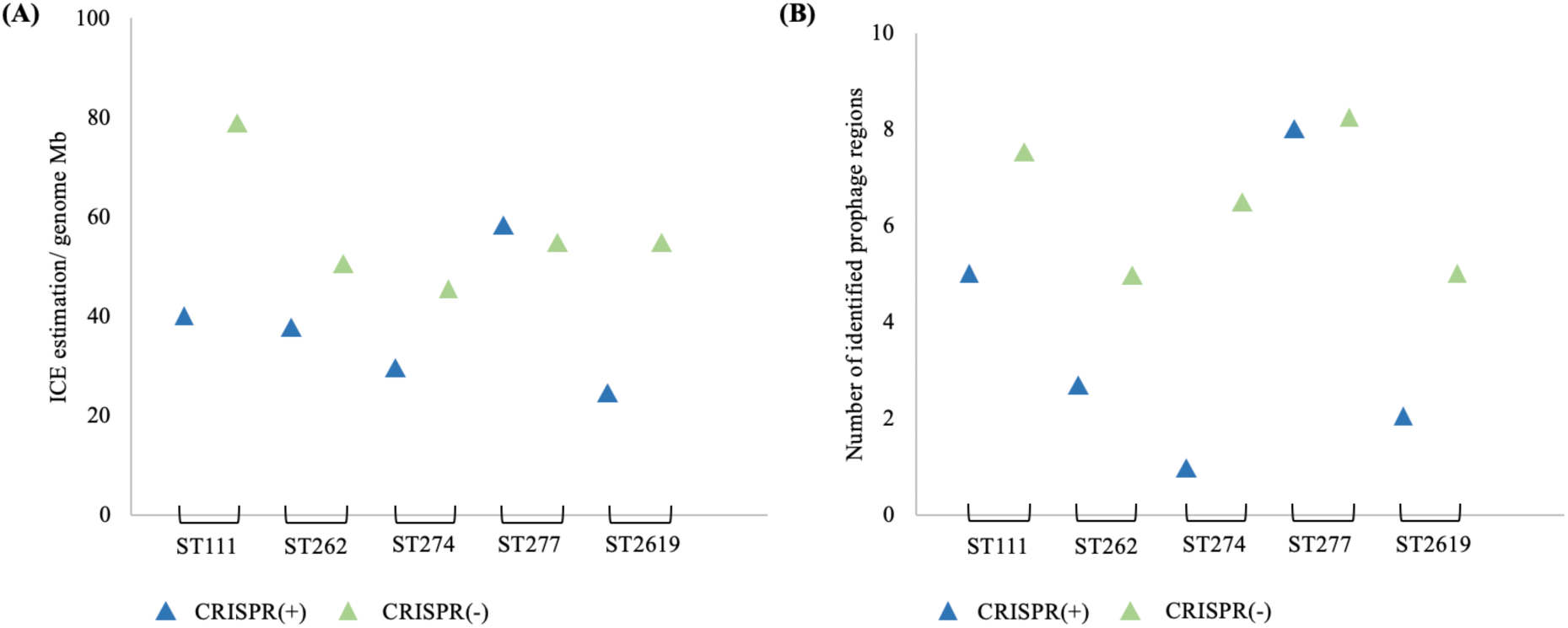
Comparative abundance of ICE and prophage regions in closely related CRISPR(+) and CRISPR(-). Panel A shows the abundance of ICE genes (/genome Mb). CRISPR(-) genomes contain a higher abundance of ICE genes compared to their CRISPR(+) counterparts (paired sample t-test p < 0.05). Panel B shows the identification of prophage regions. CRISPR(-) genomes contain a higher number of identified prophage regions compared to their CRISPR(+) counterparts (paired sample t-test p < 0.05).

### Spacers in CRISPR(+) genomes map to mobile elements present in closely related CRISPR(-) genomes

If CRISPR-Cas constrains HGT, then the spacers found in isolates with a functional CRISPR-Cas system should match to mobile elements that are present in the genomes of closely related isolates that lack CRISPR-Cas. To test this hypothesis, we aligned CRISPR(+) and CRISPR(-) complete genome representatives in ST111, ST262, ST277 and ST2619 (Fig. 5), and extracted regions unique to CRISPR(-) genomes. Crucially, we found that 10%-20% of spacers from the CRISPR(+) genomes matched targets unique to the CRISPR(-) genomes (Table 2). Matching spacers had a diversity of targets, including both phage and conjugative genes (Table 2). For example, ST111 CRISPR(-) genomes are rich in ICE (Fig. 6A), and ST111 CRISPR(+) genomes carry spacers that target conjugative transfer machinery genes (Table 2). Two spacers mapped to regions annotated as intergenic, however it is possible that these regions have been missed by genome annotation or had degenerate coding sequences(17). The identification of CRISPR(+) spacers that map to mobile elements in their closely related CRISPR(-) counterparts provides compelling evidence that CRISPR(+) systems prevent the acquisition of these elements. It also provides promising evidence that an absence of CRISPR inhibition of HGT has contributed to genome divergence and expansion in these lineages. To our knowledge, this is the first analysis of CRISPR-Cas in relation to intra-ST *P. aeruginosa* genome evolution in such a way.

## Conclusions

Broad scale comparisons across the diversity of *P. aeruginosa* reveal that CRISPR-Cas systems are associated with smaller genome size (Fig. 1) and a higher GC content (Fig. 2), which is indicative of reduced acquisition of low GC mobile elements. Anti-CRISPR genes negate the effects of CRISPR-Cas systems on genome size (Fig. 1), suggesting that Acr genes provide effective long-term inhibition of CRISPR-Cas function. To gain insights into how CRISPR-Cas systems restrict HGT, we focused on comparing the genomes of closely related strains that are CRISPR(+) or CRISPR(-). Phage are a main target of *P. aeruginosa* spacers (Fig. 3), and CRISPR-Cas systems constrain the acquisition of prophage (Fig. 6B). Most isolates with a functional CRISPR-Cas system carry spacers that target either ICE or the conserved conjugative transfer apparatus used by ICE and conjugative plasmids (Fig. 4), and CRISPR-Cas systems also constrain the acquisition of ICE (Fig. 6A). These comparisons between closely related isolates provide clear-cut evidence for the effects of CRISPR-Cas on genome composition, and they provide an important complement to the broad-scale analyses that focus on the association between CRISPR-Cas and HGT at larger phylogenetic scales (18, 28) and experimental studies that investigate the influence of CRISPR-Cas on the transfer of individual elements (18, 19, 85). Collectively, our results provide further evidence to support the hypothesis that CRISPR-Cas can act as an important constraint on HGT in bacteria (10, 18, 28), and *P. aeruginosa* is an example of an important bacterial pathogen where this seems to be the case (18, 28).

Although CRISPR-Cas systems were initially characterized as phage defence mechanisms (9, 10), it is becoming increasingly clear that CRISPR-Cas systems also target mobile genetic elements (19-21, 86). Why does CRISPR-Cas target these elements? On the one hand, the tight correlation between the number of spacers targeting phage and total spacer count suggests that spacers targeting mobile genetic elements may simply be acquired as a non-selected by-product of CRISPR-Cas systems that have been selected to rapidly acquire spacers that target invading phage. Alternatively, it is possible that selection favours the acquisition of spacers that target mobile elements due to the costs associated with these elements. It is compelling that we identified a large number of spacers that target the conjugative transfer apparatus, suggesting that selection has favoured the acquisition of these spacers, perhaps as a result of fitness costs (87) or increased phage susceptibility (84, 88) associated with the expression of conjugative machinery. *P. aeruginosa* uses the type VI secretion system to attack neighbouring cells in response to contact with conjugative pili (89), and CRISPR-Cas targeting of conjugative pilus may prevent *P. aeruginosa* from acquiring elements that trigger attacks from their kin. In line with our findings, experimental work has shown CRISPR-Cas systems in *S. epidermidis* can be effective at preventing plasmid transmission (21), and Acr genes have recently been identified on plasmids that can help overcome this immunity (90). The ICE database that we queried is likely an underrepresentation of the true diversity of ICE that can be transferred to *P. aeruginosa*.

Plasmids are widespread in bacteria, and they play a key role in HGT (91, 92). At a broad scale, plasmid carriage is higher in CRISPR(-) genomes than in CRISPR(+) counterparts (19), suggesting that CRISPR systems play an important role in constraining the transfer of plasmids between bacteria. Plasmids have played an important role in acquisition of resistance genes in *P. aeruginosa*, including carbapenemases (93-97), highlighting the importance of understanding constraints to plasmid transfer in this pathogen. Systematic surveys of the abundance of plasmids in *P. aeruginosa* are lacking, in part due to the challenges of identifying plasmids from short read assemblies, but it is clear that plasmids are much less abundant than ICE, as *P. aeruginosa* genomes typically contain multiple ICE (Fig. 5). Interestingly, many *P. aeruginosa* plasmids lack conjugative genes (46), suggesting that conjugative plasmids may be restricted to a sub-set of the diversity of *P. aeruginosa* (97) or carry genes that overcome CRISPR-Cas immunity.

Our search for spacers that target mobile elements was based on searching databases that represent known ICE, phage and plasmids. These databases clearly under-represent the diversity of mobile elements that can be transferred to *Pseudomonas* (17). Given this, alongside the stringent search parameters used to avoid false positive discovery, our study provides a conservative estimate of the limitations to HGT created by CRISPR-Cas immunity. Specifically, we speculate that many of the *P. aeruginosa* accessory genes targeted by spacers (this accounts for ∼1/3 of spacers) may be uncharacterised cargo genes carried by ICE or plasmids.

CRISPR-Cas systems are present in an estimated ∼85-90% of archaea, but are significantly less prevalent in bacteria (1). The molecular mechanisms of CRISPR-Cas are clear (4, 5), but studies have produced conflicting results on the importance of CRISPR-Cas to genome evolution (18, 21, 23-27). This study contributes to a growing body of evidence showing that CRISPR-Cas systems can constrain HGT (18, 21, 27, 28, 90). HGT plays a key role in adaptation to new niches and stressors (84, 98), suggesting that the xenogenic immunity provided by CRISPR-Cas may ultimately constrain in the ability of bacteria to adapt to new challenges. For example, it is possible that the large-scale use of antibiotics in agriculture and medicine has provided microbes lacking CRISPR-Cas with an important advantage due to their enhanced ability to acquire mobile resistance genes (18). Horizonal transfer plays a key role in the maintenance of mobile genetic elements (99, 100), and conjugation provides a simple and efficient mechanism for horizontal transfer. A second challenge for future work will be to understand how mobile elements overcome the obstacles associated with CRISPR mediated immunity. Recent work has shown that *Acr* genes allow plasmids to transfer in the face of CRISPR-based immunity (90), and it is possible that mobile elements have evolved the ability to exploit other mechanisms of horizontal transfer as a response to CRISPR-based immunity, such as transduction, outer membrane vesicles, and DNA uptake.

## Acknowledgements

This project was supported by Wellcome Trust Grant 106918/Z/15/Z held by RCM. We would like to thank Mike Brockhurst, Alvaró San Millan, and reviewers for comments on the manuscript.

